# Species coexistence and overlapping distributions in a metacommunity are compatible with niche differences and competition at a local scale

**DOI:** 10.1101/2020.07.10.196758

**Authors:** Maxime Dubart, Patrice David, Frida Ben-Ami, Christoph R. Haag, V. Ilmari Pajunen, Dieter Ebert

## Abstract

Niche partitioning is the most studied factor structuring communities of competing species. In fragmented landscapes, however, a paradox can exist: different taxa may competitively dominate different types of habitat patches, resulting in a form of spatial niche partitioning, yet differences in long-term distributions among species can appear surprisingly small. This paradox is illustrated by an emblematic metacommunity - that of *Daphnia* spp. in rockpools on the Finnish Baltic coast, where three species compete with each other, have distinct ecological preferences, yet largely overlap in long-term distributions. Here we examine how metacommunity models that explicitly estimate species-specific demographic parameters can solve the apparent paradox. Our research confirms previous studies that local extinction rates are influenced by environmental variables in a strong and species-specific way and are considerably increased by interspecific competition. Yet, our simulations show that this situation exists alongside interspecific differences in realized niches that are, overall, small, and identified three main explanations for this compatibility. Our results illustrate how state-space modelling can clarify complex metacommunity dynamics and explain why local competition and niche differentiation do not always scale up to the landscape level.

## Introduction

Understanding the determinants of community composition and turnover is a long-standing challenge in ecology and is mandatory for explaining species distributions in time and space. Two classes of models have addressed this topic: models of niche differentiation, which allow species coexistence by partitioning the niche space (Hutchinson 1959, Chase and Leibold 2003), and neutral models (MacArthur and Wilson 1967, Bell 2001, Hubbell 2001), where stochastic processes (e.g., birth, death) drive the dynamics, and species coexistence is transient, resulting ultimately in the extinction of all but one species unless counteracted by speciation or migration. These models represent extremes along a continuum from fully deterministic (niche model) to fully stochastic community assemblies (Gravel et al. 2006). In the deterministic (niche) models, species presence, and therefore coexistence or exclusion depends on the interplay of both abiotic (*i*.*e*. environmental requirement) and biotic (*i*.*e*. interspecific interactions) conditions. In purely neutral models, species relative abundances follow a random walk as species are considered ecologically equivalent, and long-term community composition is not predictable.

Through migration, local communities are linked in a larger metacommunity (Leibold et al. 2004) where species diversity and coexistence can be viewed at different scales (see e.g. Amarasekare, 2003). On a local scale, niche and neutral models may explain species presence, while on the metapopulation scale, species diversity may depend on environmental heterogeneity, migration among communities and differentiation among species. Identifying mechanisms and quantifying their relative importance for community assembly and species coexistence is a difficult task in natural systems (Gilbert and Bennett 2010, Smith and Lundholm 2010, Tuomisto et al. 2012).

In a metacommunity context, a specie’s direct interactions with its biotic and abiotic environment may not sufficiently describe its niche. Indeed, each species faces a complex mosaic of favorable environments that allow it to maintain itself over the long term and unfavorable ones that do not. At any given time, a species may be absent from a favorable habitat (e.g., due to stochastic extinction, competitive exclusion, or dispersal limitation) and present in unfavorable ones (e.g., due to random colonization, recurrent immigration or positive interspecific interactions). Therefore, characterizing the fundamental and realized niche of a species requires teasing apart the roles of environment, interspecific interactions, and dispersal; for this, spatio-temporal data that records community composition over multiple locations and points in time is necessary (Fidino et al. 2019).

The Levins metapopulation model (Levins 1969) provides a framework for understanding how colonization and extinction rates in habitat patches influence the way species occupy fragmented space. This metapopulation model has been extended to include patch heterogeneity, for example in the Incidence Function Model (Hanski 1994), and to include multiple species and their interactions (Ilkka Hanski, 1983; Slatkin, 1974; see Miller et al. 2012, Davis et al. 2017, Fidino et al. 2019, for empirical examples). A state-space approach has also been developed to deal with imperfect detection (MacKenzie et al. 2003). With spatio-temporal data, multi-species metapopulation models can now, thus, disentangle the determinants of community composition, although, to date, these state-space models have mainly been applied to single species (Royle and Kéry 2007, Martin et al. 2010, Eaton et al. 2014) and only rarely to metacommunities (Miller et al. 2012, Davis et al. 2017). State-space models offer advantages for analyzing metacommunities, though: they allow us to quantify colonization and extinction rates depending on both environment and interspecific interactions; they can also consider dispersal explicitly, and they can accommodate imperfect data (MacKenzie et al. 2002, Guillera-Arroita 2017).

This study extends the approach of Dubart et al. (2019), who used multispecies metapopulation models to estimate colonization and extinctions rates based on environmental conditions and the presence of other species, while also accounting for imperfect species detection probabilities and non-equilibrium situations. Here, we extend this approach by including space explicitly and reformulating the initial model as a continuous time version, so that it applies to a wider range of real metacommunities. We apply it to analyze a long-term dataset of three *Daphnia* species that form a rockpool metacommunity.

### The study system

The planktonic crustaceans of the genus *Daphnia* that inhabit the rockpools of the Baltic Skerry islands have been studied for over a hundred years (Levander 1900, Järnefelt 1940, Lagerspetz 1955). Three species, *D. magna, D. longispina* and *D. pulex*, are typically found in these rockpools, but at any time, less than 50 % of the pools are occupied by any one species, and local extinction rates are about 20 % per year, indicating extremely dynamic metapopulations (Pajunen 1986, Pajunen and Pajunen 2003, Ebert et al. 2013). Attempts to define their ecological niches have shown a large overlap in the niches of the three *Daphnia* species (Ranta 1979, Pajunen and Pajunen 2007), suggesting that competition is a strong factor shaping presence/absence patterns, as experiments have shown (Hanski and Ranta 1983, Bengtsson 1989, 1991, 1993). Although environmental factors such as temperature, water chemistry, food, and predation have been suggested as possible mediators of competition, no general patterns have been identified (Bengtsson 1986, Ranta and Espo 1989, Milbrink and Bengtsson 1991, Lehto and Haag 2010). Several studies have attempted, albeit without consensus, to determine the driving principles underlying this system’s pronounced dynamics: Hanski & Ranta (1983) considered all rockpools as equal and suggested that a trade-off exists between competitive ability and dispersal propensity (i.e. competition-colonization tradeoff). Pajunen (1986) and Pajunen & Pajunen (2003) have suggested a continent–island model, while Hugueny et al. (2007) favored a Levin-type metapopulation model, with all ponds being equal.

The continent–island view was based on the finding that some rockpools contain more stable populations than others and may therefore contribute more to the pool of migrants. However, Altermatt and Ebert (2010) suggest an inverse continent–island model, pointing out that because small, unstable rockpools are more abundant, they contribute most to the migrant pool. Furthermore, experimental tests have shown that not all pools provide equally suitable habitats, suggesting that colonization rates depend on pool quality (Ebert et al. 2013). Evidence also suggests that habitat preferences may differ among species (Lehto and Haag 2010), although generally little is known about the species-specific spatio-temporal variation in habitat suitability. All in all, although research on the *Daphnia* rockpool metapopulation has identified some mechanisms that contribute to its pronounced dynamics, the relative importance of the different factors remains unknown, and despite all these years of research, the coexistence of these species is still enigmatic. In this study, we investigate the contributions of spatial structure, biotic and abiotic environmental factors, and species interactions in driving extinction and colonization dynamics in this metacommunity, as well as occupancy patterns. We use the above-mentioned metacommunity model to analyze 35-year time series data on three Daphnia species from 546 rock pools on 17 islands in southern Finland.

## Material & Methods

*Daphnia* are planktonic crustacean that inhabit standing fresh and brackish water bodies worldwide. Here, we study three species, *D. magna* Straus, *D. pulex* Leydig, and *D. longispina* O.F.Müller, that co-occur in rockpools of the Baltic sea and Atlantic coasts in Fennoscandia, where they form large metapopulations. Specifically, we studied all rockpools (*n* = 546 in total) in an area of ∼1.5 km^2^ near Tvärminne Zoological Station (coordinates: 59.844, 23.249) in southern Finland, an area that comprises a total of 17 Skerry islands. The rockpools are situated along the shore of the Baltic Sea, keeping them clear of plant overgrowth and are mostly filled by rainwater, although the water chemistry of pools closer to the shore is also affected by wave action and sea water spray. During winter, the ponds may freeze solid, and, in summer, they may dry up completely. Pool volumes range from about 3 to 50000 L with a median of about 200 L (Altermatt et al. 2009).

### Data collection

For this study, we used time series of presence/absence data collected twice a year from August 1982 to August 2017. Rockpools were visited every year in late May/early June (spring sample) and in late July/August (summer sample), usually after rainy periods, but long enough after the rain to allow *Daphnia* to hatch from resting eggs (Pajunen 1986, Pajunen and Pajunen 2003). Using a handheld net (250 µm mesh size), we sampled the free water of each pool, inverted the net into a white bowl and checked for the presence of *Daphnia*. If they were present, a small sample was taken to the laboratory. If different phenotypes were visible, care was taken to include these. In the laboratory, samples were checked with a stereomicroscope to identify the species. This procedure suffers from two potential sources of error. First, *Daphnia* may be present but overlooked by the observer, for example, if they are present in very low density, or if one species is much more rare than a co-occurring one. It may also happen if *Daphnia* are present only as resting eggs (“ephippia”). which are highly resistant to a wide range of conditions, including freezing and drought (Hanski and Ranta 1983, Altermatt and Ebert 2008), and which cannot be systematically sampled. In each of these cases, the error leads to a false absence record. Second, a species may also be misidentified (or misrecorded), which results in a false presence as well as a false absence record. In addition to these two possible sources of errors, ponds that had dried up during the recording period presented uncertainty, as the presence of resting eggs could not typically be verified. In these cases, the presence/absence of each species was recorded as “not assessed”. To estimate the repeatability of our data, seven surveys between 2009 and 2012 were repeated a second time by a different team of researchers within two weeks of the first survey. These repetitions allowed us to assess the likelihood of missing a given *Daphnia* species (i.e. the detection probability) and the frequency of misidentification. As the latter (the apparent replacement of one species by another within two weeks, which we attribute to misrecording or misidentification) occurred only once across all seven repeated surveys, we discounted misidentification and misrecording from further analyses.

Following earlier protocols (Pajunen 1986, Ebert et al. 2001, Pajunen and Pajunen 2003, 2007, Altermatt and Ebert 2008, Altermatt et al. 2009, Altermatt and Ebert 2010), GPS coordinates (using a Garmin GPSMAP 76CS), island ID and a number of environmental variables were recorded for each pool. As opposed to earlier studies, we included all pools in the study area, including those with no history of *Daphnia*, such as pools very close to the sea, so that no inclusion-criteria had to be used. Distance to the sea and height above sea level were measured. Pool surface area was estimated by multiplying its longest axis by widest width (measured perpendicular to the longest axis) at maximal water level divided by 2. Maximal pool depth was measured. Catchment area was estimated as described by Altermatt et al. 2009. During several August surveys, we also estimated the percentage of the surface water covered by terrestrial plants using the categories: 0, >0-5, -25, -50, -75, -95, >95 % of the water surface. We also measured water conductivity and pH using a hand-held portable pH meter and conductivity device several times across multiple years. For plant coverage, conductivity and pH, we calculated the mean and the standard deviation across all measures. The Baltic sea has no tides, but the sea level varies depending on wind, atmospheric pressure and other factors. Rockpools close to the shore may alternate between being submerged and then being above sea level for extended periods, long enough for *Daphnia* to colonize and produce a population. Pools below the maximal sea level are labelled as “sometime submerged” (binary variable). *D. magna*, but not *D. longispina* nor *D. pulex*, can survive in brackish water up to or even greater than the salinity of the Baltic Sea, though none of the species occurs in the sea.

The ecological variables and GPS coordinates are complete in our dataset, but the presence/absence data are not: individual data points are missing from when pools dried up, which especially affected the smaller, shallower pools with smaller catchment areas (Altermatt et al. 2009). The dataset has also missing values from spring 1993 when the survey could not be conducted. For most of the rockpools, the final dataset includes time series of 70 samples of present/absence data for three *Daphnia* species (nearly 40,000 total pool visits).

The ecological variables for each pool (Table S1) were summarized using a principle component analysis. The first two principle components together explained 43.9% of the variance (first axis: 25.4%, second axis: 18.5%) and were ecologically meaningful, as they describe the impact of the marine habitat (e.g., closeness to the sea, water chemistry, plant cover, horizontal axis on Fig. S1), and the effect of geo-physical features independent of the sea (e.g. pool size, depth, catchment area, vertical axis on Fig. S1), respectively. We use the terms “marine to terrestrial” to describe ecological variation among pools along the first axis and “small to large” habitats for the second axis. In analyzing extinction and colonization dynamics, we distinguished between events that happened between the spring and the summer survey (“summer events”) and events that happened between the summer survey and next year&s spring survey (“winter events”).

### Model

In order to estimate *Daphnia* metapopulation dynamics, we used a state-space modeling approach (MacKenzie et al. 2003, Royle and Kéry 2007) based on one developed by Dubart et al. (2019), which models colonization and extinction rates as a function of the environment and the presence of other species, while accounting for imperfect species detection. This model allowed us to consider colonization rate as dependent on overall metapopulation occupancy, as in Levins’ metapopulation model (Levins 1969). In contrast to Dubart et al (2019), who modeled extinction and colonization as discrete events, we used transition probabilities from a continuous time model to account for the extremely dynamic nature of the *Daphnia* metapopulations (Pajunen 1986, Pajunen and Pajunen 2003). Furthermore, we extended the previous model to account explicitly for space, given the distance-dependent dispersal seen in the *Daphnia* metapopulations (Haag et al. 2005, 2006).

The idea behind state-space modeling approaches is to distinguish between the true state and the observed state (MacKenzie et al. 2002), hence accounting for imperfect species detection. The true state (*x*_*S,i,t*_, for species *s*, in site *i* at time *t*), and the observed state (*y*_*S,i,t*_) are linked as follows:

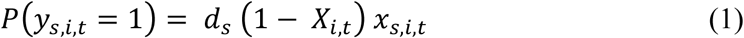

where *X*_*i,t*_ is an indicator variable that equals 1 for a site that was dry at the time of sampling (species cannot be detected in dry sites), and 0 otherwise, and *d*_*S*_is the detection probability for species *s* in a non-dry site - *d* = *P*(*y*_*S,i,t*_ = 1 | *x*_*S,i,t*_ = 1). The latter can be jointly estimated with all other model parameters (see below) and is primarily informed by the repeated surveys.

Next, we modeled the transitions among states (from *x*_*S,i,t*_ to *x*_*S,i,t*+1_) as a function of species-specific and site-specific colonization (*γ*_*S,i,t*_) and extinction (*e*_*S,i,t*_) rates: for a given species in a given site, this gave the following probabilities of presence at time *t*+1 (depending on the presence or absence at time *t*, species and site subscripts were dropped for readability):

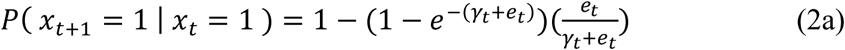

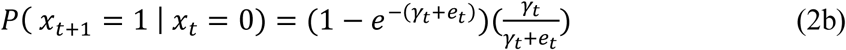

The two probabilities of species absence (*P*(*x*_*t*+1_ = 0 | *x*_*t*_ = 1) and *P*(*x*_*t*+1_ = 0 | *x*_*t*_ = 0)), are given by the complements of equations (2a) and (2b), respectively.

Taking Eq. 2b as an example, the probability of having *x*_*t*+1_ = 1 knowing *x*_*t*_ = 0 at a given site (i.e. the probability that a species is present at time *t+1* knowing it was absent at time *t*) is the product of the probability of having at least one event in the time step 1 – *e*^−(*γ*+*e*)^ and the probability that the last event, if there is one, is a colonization 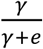 (note that *e*^−(*γ*+*e*)^ is the Poisson probability of no event with rate *λ = γ* + *e*).

The colonization rate (*γ*_*S,i,t*_) depends on the per-capita colonization rate (*c*_*S,i,t*_) and on metapopulation occupancy (*p*_*S,i,t*_): *γ*_*S,i,t*_ = *c*_*S,i,t*_ *p*_*S,i,t*_. Per-capita colonization and extinction rates are species and site specific and depend on environmental variables and species interactions:

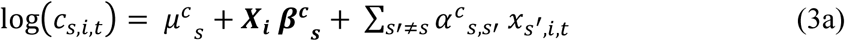

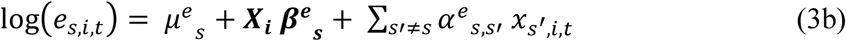

Where *μ*^*e*^and *μ*^*c*^ represent extinction and colonization rates, respectively, in a site with average characteristics and no other species present, ***X***_***i***_ is a row vector of environmental variables (standardized to zero mean and unit variances), and ***β***^***e***^_***S***_ and ***β***^***c***^_***S***_ are the species-specific regression coefficients for extinction and colonization, reflecting that these parameters depend on environmental variables. The parameters 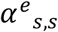 and 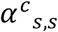 measure competitive (or facilitative) effects of species *s’* on extinction and colonization rates, respectively. Note that 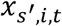 are latent variables (i.e. estimated true state) rather than observed states. Since 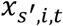 are indicator variables (equal zero or one), *μ*’s represent rates in a site with average characteristics (***X***_***i***_ = **0**) and no competitors.

Following Levins (1969), colonization rate (*γ*) depends on overall metapopulation occupancy (*p*) as propagules pressure is expected to increase with the number of potential source populations. However, when dispersal is limited (in terms of distance), the sources from which colonists originate may vary depending on the focal site. We therefore use a site-specific metapopulation occupancy (*p*_*S,i,t*_) defined as follows:

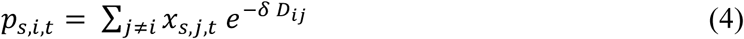

where *D*_*ij*_ is the Euclidean distance between sites *i* and *j* (in meters), and 1/*δ* the mean colonization distance. In this way, the contribution of a given site to colonizing a focal site decreases exponentially with the distance between sites. Note that we therefore assume symmetrical connectivity patterns. Using this formulation, our *c* does not have the same meaning (corresponding to per-capita colonization) as Levins’ *c*. We therefore scale *p*_*S,i,t*_ to recover an analog of Levins’ *c* (see Appendix. A for derivation), so that the final *p*_*S,i,t*_ reads:

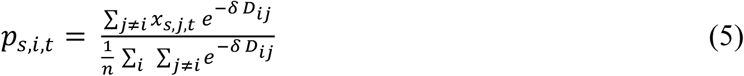

As we only modeled transitions, we made no assumptions about the (quasi-)equilibrium state, nor about initial spatial distributions or abundances of species. The initial state (initial spatial distributions and abundances) was estimated using an additional species-specific parameter (*ψ*_*S*_).

### Implementing the model

We split the dataset into five groups of islands with large distances between them (>200m), so that the effect of between-group dispersal could be neglected. Our main motive was to reduce computation time, as only submatrices needed to be considered for the calculations of *p*_*S,i,t*_ and previous studies have found that colonization events beyond the 200-m threshold are rare (Pajunen 1986, Pajunen and Pajunen 2003), although it is possible to find distances over 200 meters within islands groups. Island groups were considered as independent for parameter estimation, but parameters are shared by the five groups and are estimated simultaneously.

We relied on a Bayesian approach to estimate model parameters and implemented the model using JAGS (Just Another Gibbs Sampler, Plummer 2003) through its R interfaces ({rjags} and {coda} R packages, Plummer et al. 2006). We used wide prior distributions for all parameters: ***μ***’s, ***β***’s, and ***α***’s ∼ *N*(0,0.1) (0.1 being a precision parameter - the inverse of the variance), detection probabilities (***d***) and initial probabilities of occupancy (***ψ***) followed a uniform distribution on [0;1], and the exponential decay of the dispersal kernel ***δ*** ∼ *U*(10^−5^, 0.5). We ran six chains, with random initial values for each of these, for 20,000 iterations, with the first 10,000 iterations discarded as burn-in periods. We assessed convergence visually and used several diagnostic tools from the {ggmcmc} R-package (Fernández-i-Marín 2016). The potential scale reduction factors 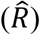 were computed following Gelman et al. (2013) and were all less than 1.03 (mean = 1.0036, *CI*95 = [1; 1.02]).

### Investigating species-environment relationships

Environmental effects estimated by the model represent effects without competition and does not include dispersal effects or habitat heterogeneity. It follows that a pond embedded in the full habitat matrix could harbor a population for some time due to colonization from surrounding, more favorable habitats, but could not support a population over the long term if it were surrounded by habitats similar to itself. Indeed, in the absence of competition, the long-term presence probability at site *i* is 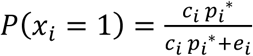, with *p*_*i*_^∗^ being long-term occupancy in the surrounding sites - the expectation of Eq. 7. Thus, *p*_*i*_^∗^ encapsulates both dispersal effects and habitat heterogeneity but is not easily derived analytically. It follows that defining the environmental space allowing species persistence (i.e. fundamental niche) in a metapopulation relies on simulating metapopulations with a single habitat type common to all patches, excluding heterogeneity. The question therefore becomes whether - knowing the spatial arrangement of sites and the dispersal ability of the species - a given habitat type (a set of values of environmental variables) allows species persistence at the landscape scale? Defining the realized niche (i.e. accounting for species interactions) is based on similar arguments but *P*(*x*_*i*_ = 1) takes a more complicated form *E*[*c*_*i*_], *E*[*e*_*i*_] and *E*[*p*_*i*_^∗^] depend on the presence probability of other species. As there are no analytical expressions for these quantities, we rely on simulations based on parameter estimates.

The simulation model for the full landscape of 546 pools and their explicit spatial arrangement and species dynamics follows Eqs. 2. Initial occupancy was set to 0.2 (the approximate mean over all species) for all species, and the spatial distribution of initially occupied sites was randomly drawn. We removed habitat heterogeneity by setting all site covariate values to those observed for one of the 546 sites, then explored all 546 environmental variable combinations in turn, setting competition to zero or not. These 546 combinations fall into three categories based on whether the metapopulation (i) persists in a uniform landscape even in competition with other species, (ii) persists only in the absence of competition, or (iii) does not persist. Category (i) is the metapopulation estimate of the realized niche, whereas the fundamental niche is defined by (i) and (ii) together. Each simulation was run for 500 cycles (i.e. years) and replicated 500 times, each time with a new set of parameters drawn from the full joint posterior distribution provided by the statistical model (thus also including model uncertainty). A metapopulation was considered to have persisted if the final occupancy was above zero in at least 50 % of the replicates (results stay qualitatively similar when considering other thresholds, see Fig. S2).

Finally, we simulated species dynamic in the heterogeneous, “real” landscape (i.e. each site with its own combination of covariables) with three aims: (i) to assess the fit to data by comparing the observed 35-yr series to the simulated 35-yr series with the same initial situation (short-term simulations), (ii) to estimate the long-term stationary distribution of occupancy for each species (long-term simulations), and (iii) to assess how dispersal limitation and interspecific interactions influenced long-term and short-term occupancy. To this end, we performed a set of simulations that included both the observed heterogeneity in environmental variables among sites and the explicit spatial arrangement of the sites. Simulations were initialized using either the observed distributions at the beginning of the dataset (in spring 1983 sample) or a random drawn of occupied sites with probability 0.2. Simulations were run for a time span of 2,500 years to get the quasi-stationary distribution (long-term simulations). The first 35 years were also recorded for short-term predictions. For the long-term simulations, we visually verified that the distributions of occupancy in non-extinct metapopulations converged at the 2,500th year. Note that the final state of any isolated, finite stochastic metapopulation model is always extinction, but the non-extinct states converge to a quasi-stationary distribution before that. In a spatialized model, this may occur for an isolated subpart of the system, which we observed in some preliminary simulations, so we therefore also ran models including a small offset (*ϵ* = 5.10^−4^) to the colonization rate, mimicking an exterior colonization source, to avoid this situation. The chosen value was small enough, approximatively one external colonization every 3.5 years, to be just useful to recolonize the subpart. The initial simulation sets included the estimated dispersal kernel in the colonization function and species interactions, using all the model estimates. We also ran simulations without distance effects (i.e. using Levins’ *c* only, so that colonization would no longer be sensitive to distance but remain the same on average over all pairs of sites; we kept the offset for sake of comparability), and without interactions (setting all interspecific interaction coefficients to zero). In each case, we ran 500 replicate simulations; variation among replicates included the inherent stochasticity of the model and the uncertainty in parameter estimation (the parameters for colonization and extinction rates are redrawn for each replicate in the posterior distribution).

## Results

### Bayesian estimation of metacommunity parameters

In ponds that retained water, the estimated detection probabilities were 0.74 for *D. longispina* and *D. magna*, and 0.63 for *D. pulex* (Fig. 1A). Median colonization distances were similar for all three species (around 15-20 m.) with overlapping 95% credibility intervals (hereafter CI95) (Fig. 1B). Model intercepts estimated species-specific colonization and extinction rates in a site with average environmental characteristics (*i*.*e*. at (0,0) in the PCA space) in the absence of competing species. For all species, colonization rates during the summer were two to three times higher than in winter (see Fig. 1C). In contrast, extinctions occurred mostly during winter (Fig. 1C). Thus, the number of occupied sites tended to increase during summer and decrease during winter. The winter decrease in occupancy was most marked in *D. longispina* but also occurred in *D. magna*. It was non-significant in *D. pulex* (the CI95 of (*e* – *c*) for that species overlaps with zero).

**Figure 1.**
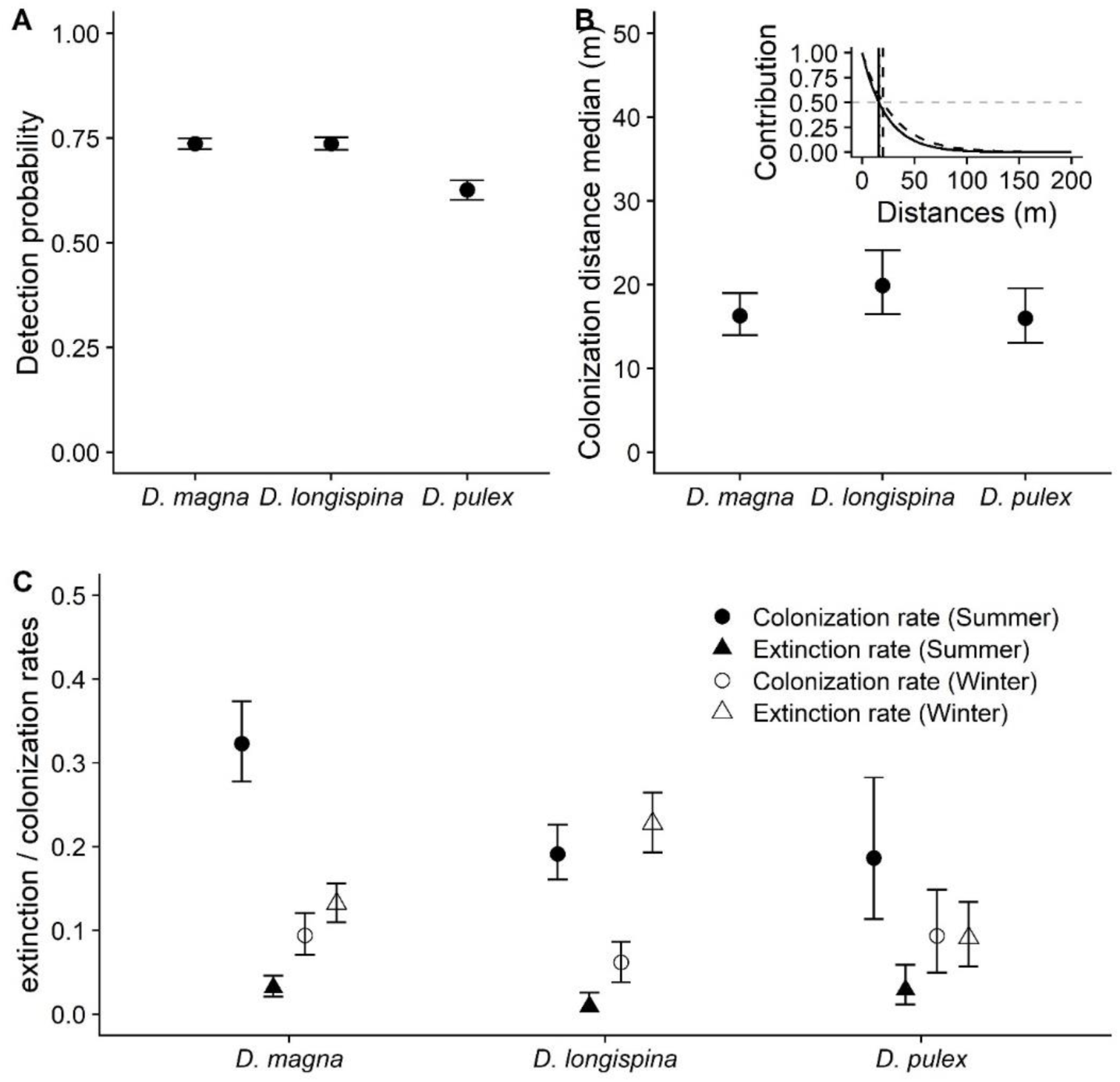
Estimated detection probabilities, colonization distance and demographic rates (median of posterior distribution with 95 % Credibility Interval CI95) for three species of Daphnia. (A) Detection probability (i.e. *P*(*y* = 1 |*x* = 1)). (B) Median colonization distances, with subpanel representing a site’s contribution to colonization as a function of the distance to the colonized site. The vertical lines provide medians (where the contribution crosses the horizontal grey dashed line, which represents 50% contribution) and are equivalent to values represented in B’s main plot. The solid lines represent both D. magna and D. pulex (not visually separable) and the dashed lines represent D. longispina. (C) Colonization and extinction rates for summer and winter for sites with average environmental characteristics (at (0,0) in the PCA space as depicted in Fig. 3) and without species interactions.

Overall, environmental variables impacted extinction rates more strongly than colonization rates (Figs. 2 and 3). On environmental PC1 and PC2, eight out of twelve CI95 for regression coefficients of colonization rates overlap with zero, versus only one out of twelve CI95 for extinction rates. The colonization rates of both *D. longispina* and *D. magna* depended slightly on PC1, but in opposite directions: In *D. longispina*, colonization rates were slightly lower in more marine pools, whereas *D. magna* colonization increased with marine influence (in winter only). Environmental variation impacted extinction rates similarly in *D. magna* and *D. longispina*, but quite differently in *D. pulex*: In *D. magna* and *D. longispina*, marine pool populations showed considerably reduced extinction rates during summer but increased extinction rates during winter (Figs. 2 and 3). In *D. pulex*, the pattern was reversed with increased extinction rates in marine habitats in summer and decreased rates in winter, though the CI for the latter overlaps with zero. In addition, extinction rates also depended on PC2. *D. longispina* and *D. magna* populations in small, shallow ponds showed increased extinction rates regardless of the season, although effects were stronger during the summer. In *D. pulex*, extinction rates increased in small, shallow ponds during the winter but, in summer, extinctions increased in large, deep pools. A first, crude criterion for site *i* suitability for a species is the estimated occupancy in Levin’s unstructured metapopulation model (1-*e*_*i*_*/c*_*i*_, averaging rates over the two seasons). According to this criterion, the same type of large, terrestrial site (see the *Expected occupancy* column in Fig. 3) appeared suitable for both *D pulex* and *D. longispina*, as both were limited by high extinctions rates in small terrestrial sites and marine sites, although these limitations arose at different times of the year (in small terrestrial sites *D. longispina* experiences more extinctions in winter, while *D. pulex* sees them in summer; the reverse is true in marine sites). Observed occupancy (averaged over years and seasons) appears to agree with our crude estimate of suitability (Fig. 3, last two columns), although more heterogeneous. Indeed, sites with low and high observed occupancy can co-occur in the same region of the environmental PC plane, which by construction, cannot happen with the expected occupancies.

**Figure 2.**
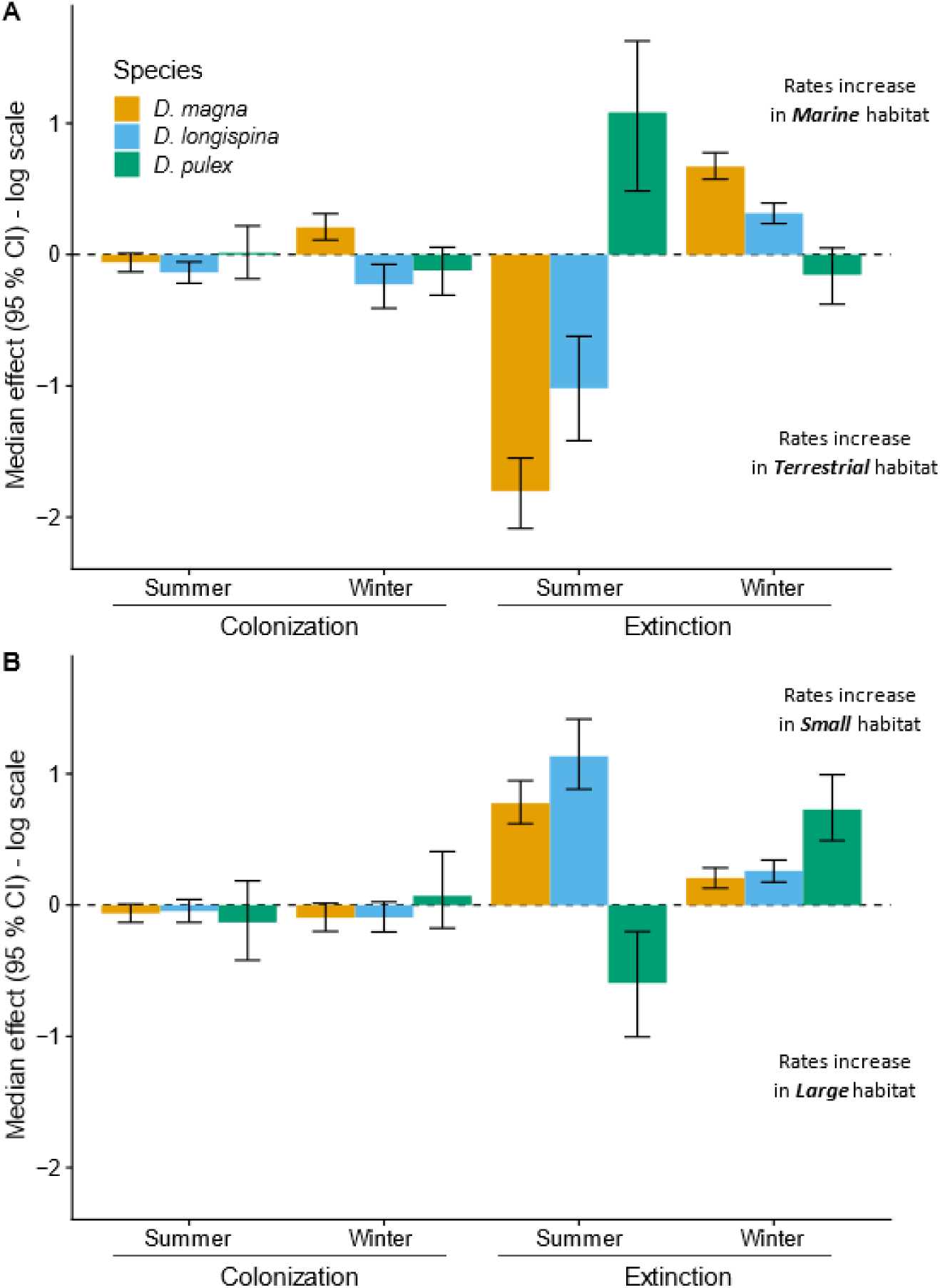
Estimated effects of environmental covariates (posterior medians and CI95) on colonization and extinction rates. (A) Log-scale changes in summer and winter demographic rates per unit of environmental PC1, mainly representing the marine–terrestrial gradient. (B) Log-scale changes in rates along the environmental gradient PC2, mainly representing geo-physical properties of the pool, such as size, depth and catchment area.

**Figure 3.**
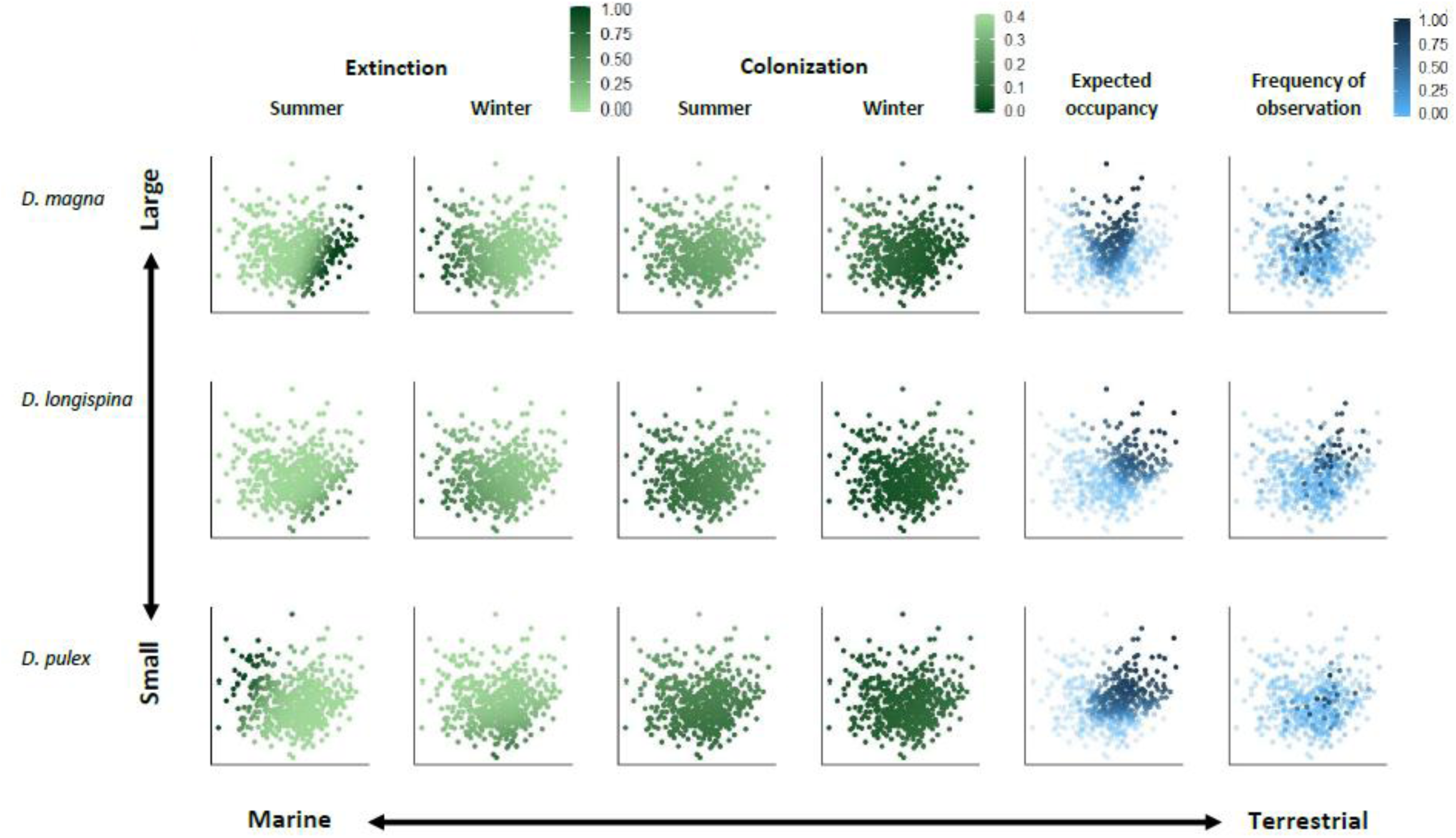
Site-specific colonization and extinction probabilities (per season), and site suitability expressed as expected occupancy in relation to environmental variables (PC axes). Colonization and extinction probabilities for site i are computed as 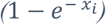, with *x*_*i*_ = *e*_*i*_ for extinction, and *x*_*i*_ = *c*_*i*_ for colonization (assuming *p*_*i*_ = 1, ∀*i*, i.e., a fully occupied metapopulation). Site-specific expected occupancy is approximated for site i as 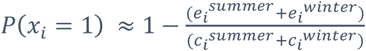, the expectation in an infinite metapopulation without environmental heterogeneity, species interactions or distance effects, i.e. the Levins model. Each row of graphs corresponds to one species (from top to bottom: D. magna, D. longispina and D. pulex), probabilities vary from 0 (light green) to 1 (dark green); colonization probabilities range from 0 (dark green) to 0.4 (light green), with light colors highlighting favorable conditions. Expected and observed occupancies, the latter averaged over seasons and years, vary from 0 (transparent light blue) to 1 (opaque dark blue).

Colonization and extinctions were both affected by the presence of other species but in roughly opposite ways (Fig. 4). Colonization rates during both summer and winter were either not significantly affected (grey in Fig. 4), or positively affected, i.e. colonization rates were higher in pools previously occupied by another species than in previously empty pools (blue in Fig. 4). The effects were generally stronger in winter, with an up to six-fold increase in colonization rates (for *D. longispina* in pools inhabited vs. uninhabited by *D. magna* the previous summer). In contrast, extinction rates tended to be affected mostly by negative interspecific interactions (i.e. increased extinction rates in ponds inhabited, in the previous survey, by other species, red in Fig. 4). Most notably, the presence of *D. pulex* and *D. longispina* in spring reciprocally increased each other’s extinction risk by over tenfold. Furthermore, *D. magna* suffered from the presence of both other species in winter, with a ∼1.7 fold increase in extinction rate, but its presence in the previous summer survey also increased the extinction risk of *D. longispina*. Only *D. magna* had a significant positive interspecific interaction, showing reduced summer extinction in pools occupied by *D. pulex* in spring.

**Figure 4.**
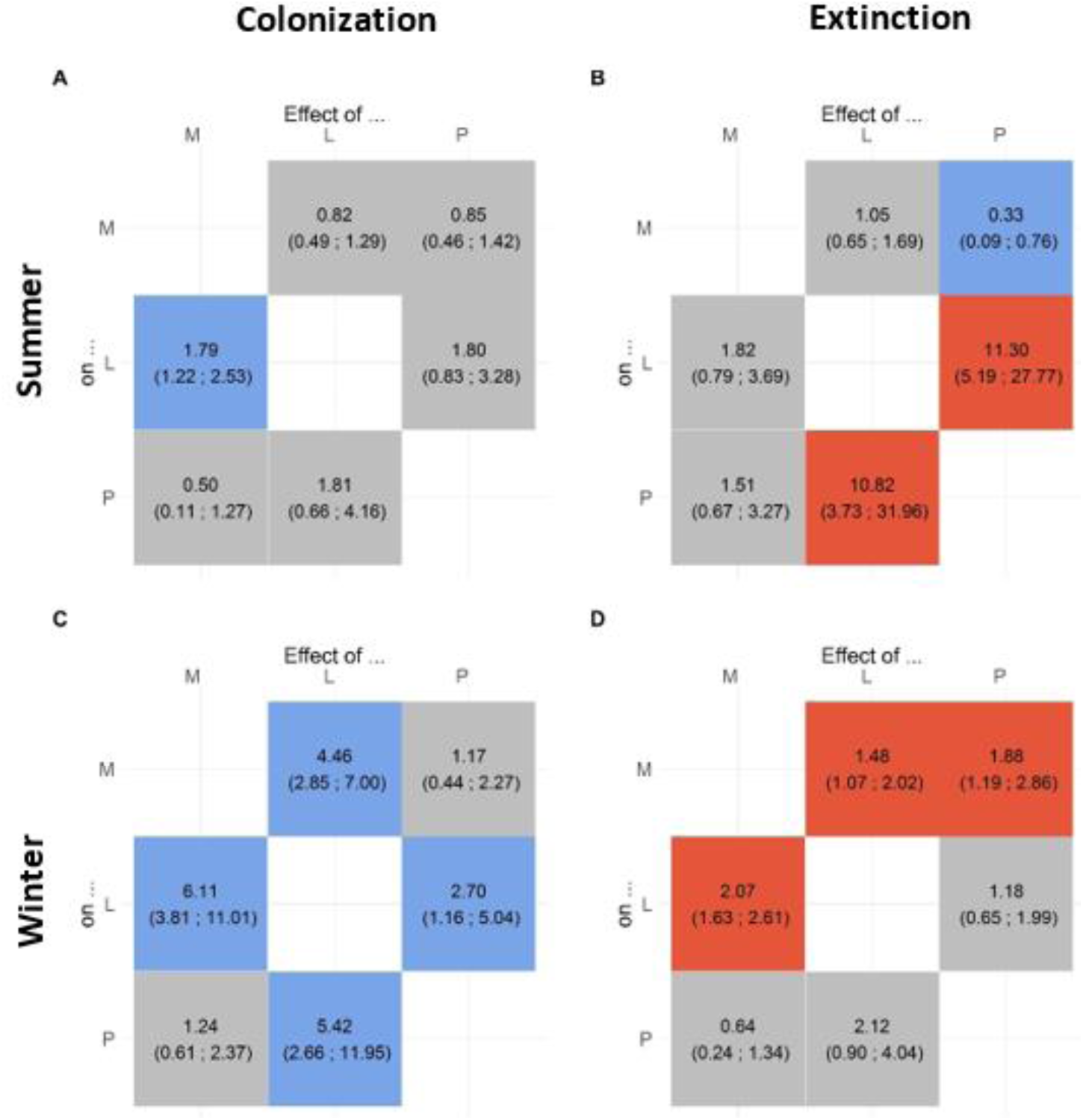
Interspecific interactions expressed as multipliers, i.e. how the presence of a species in the previous season multiplies the colonization or extinction rate of another species. Median and 95 % CI of posterior distributions are given. Coefficients with 95 % CI overlapping 1 (= no effect) are in grey. Robust competition and facilitation effects (i.e. CI not overlapping 1) are highlighted in red and blue, respectively.

### Fundamental and realized niches based on simulations of homogeneous landscapes

We previously defined the niche as the set of environmental characteristics for which a metapopulation entirely composed of sites with these characteristics would persist on the long term. In practice, we established this niche by simulating homogeneous metapopulations that preserved the number and spatial structure of the observed sites, but set them all to identical environmental values and used a 50 % threshold (i.e. the criterion for inclusion in the fundamental niche is that the simulated metapopulation is not extinct in >50% of replicate simulations at generation 500; other threshold values do not considerably change the results see Fig. S2). Based on these simulations, 63% of rockpools belonged to the fundamental niche (i.e. simulations without interspecific interactions, Fig. 5A) of *D. magna*, 52 % for *D. longispina*, and 72 % for *D. pulex*. The fundamental niches of D. *longispina* and *D. pulex* seem similar (intermediate to large-sized terrestrial sites, whereas *D. magna* preferred more marine-influenced sites. The realized niches (simulations with interspecific interactions Fig. 5B), included 64 %, 54 % and 69 % of the sites, respectively (Fig. 3). In *D longispina* and *D. pulex*, two species with very similar fundamental niches, their reciprocal impacts on each other’s summer extinction rates resulted in a divergence of their realized ecological niches: the occupancy of *D. longispina* decreased in the smallest terrestrial sites, while that of *D. pulex* decreased in the most marine ones. However, these effects were moderate, and the two realized niches still overlapped considerably. Similarly, *D. magna* showed both weak positive and weak negative effects from interacting with the other two species, and only subtle differences between fundamental and realized niches (Fig. 5C). Based on our simulations in homogeneous landscapes, the proportion of ponds with habitat types that could support a stable metapopulation of only one species (exclusive niche) were relatively low at 7% for *D. magna*, 0% for *D. longispina*, and 5% for *D. pulex (*black dots in Fig. 5D). Exclusive ponds for *D. pulex* fell into two groups: relatively small, shallow ponds with strong terrestrial or marine influence (see Fig. 5D), whereas for *D. magna*, most exclusive sites were larger, deep rockpools with a strong marine influence (Fig. 5D). Very small marine sites seemed unsuitable for all three species, and all three had largely overlapping realized niches, centered on more marine, intermediate, and terrestrial habitats for *D. magna, longispina* and *pulex* respectively (Fig. 5D).

**Figure 5.**
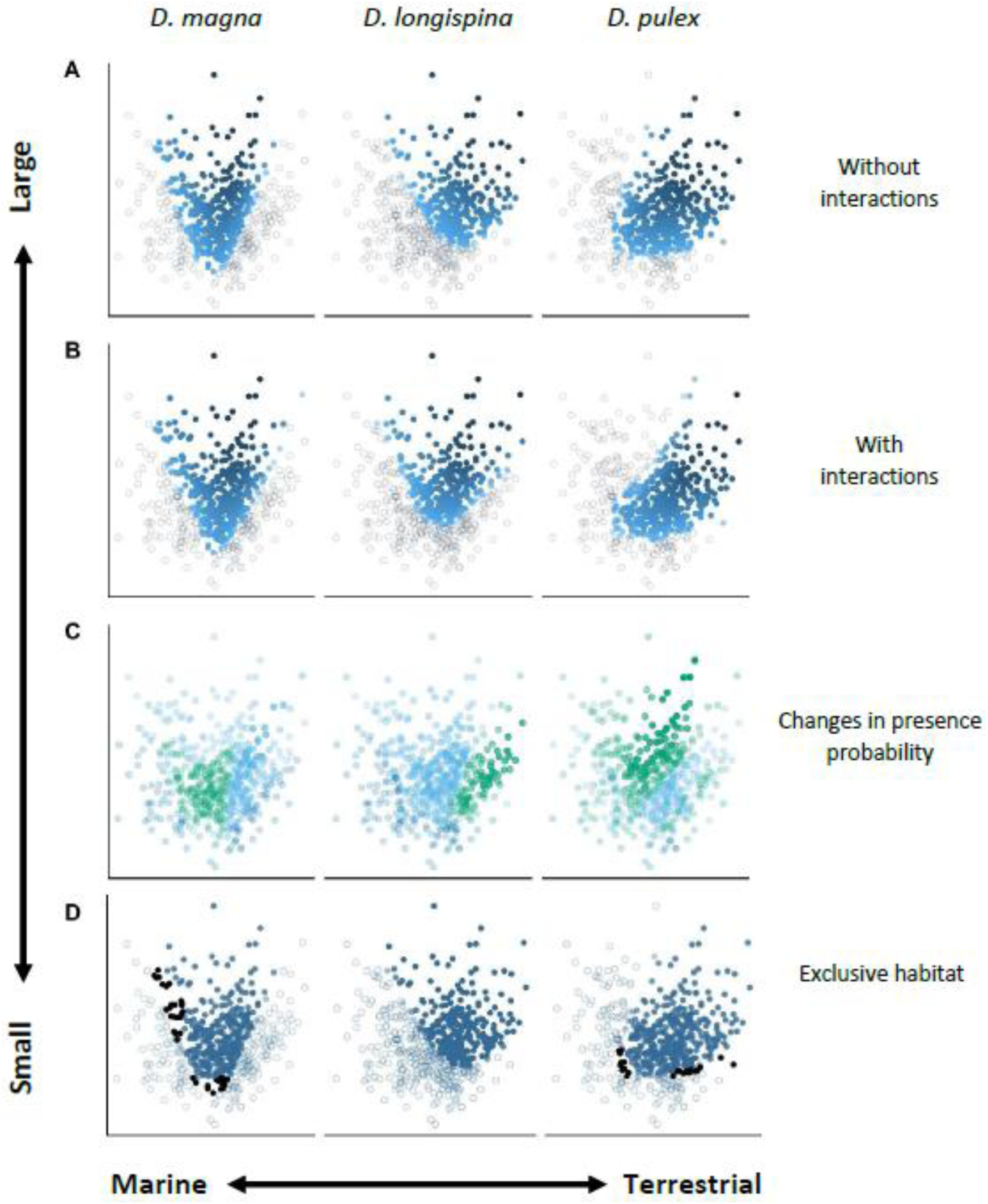
Site-specific probability of occupancy based on simulations in homogeneous environments. (A), (B) The distribution of suitable ponds within the environmental space with or without species interactions, respectively, i.e. representing the realized and fundamental niches. The light blue to dark blue gradient reflects the probability of occupancy from 0 to 1, whereas the transparency gradient represents the proportion of non-extinct metapopulations among replicates from 100 % (opaque) to 50 % (half transparent) and when < 50 %, dots are empty. (C) Shows how interactions modify site-specific probability of occupancy. Blue dots represent environments where species expected occupancies increase with interactions, and green dots, environments where they decrease. Dot transparency reflects the effect strength (defined as the difference in mean occupancy when interactions occurs or not). (D) Pools that are exclusive to each species are shown in black (i.e. exclusive niche: competition free part of the niche space), while blue dots represent remaining suitable ponds. The transparency gradient reflects the proportion of non-extinct metapopulations among replicates.

These simulations of theoretical, homogeneous landscapes predicted well the occupancies we observed in the real, heterogeneous landscape. Indeed, we saw a positive correlation between the average occupancy of a site over the 35 years of data and the simulated occupancy in a homogeneous metacommunity with the same environmental characteristics, including interspecific interactions (realized niche); this correlation was, however, stronger for *D. magna* (0.48) and *D. longispina* (0.56) than for *D. pulex* (0.23). Simulations without interactions (fundamental niche) resulted in similar correlations with the data (resp., 0.53, 0.56 and 0.20, see Table S2).

### Simulated metacommunity dynamics in a realistic, heterogeneous landscape

As expected, the offset in form of an added colonization rate from external sources, did not change results observed for the mean-field model (i.e. without dispersal limitation), indicating that our offset had little effect (compare Tables S3/S4 and S5/S6). The pattern was different with the spatialized model; indeed, the offset prevented the definitive extinction of a given species in subparts of the system. In what follows, we compared simulation results to data on two criteria (i) do they yield similar occupancies at the metapopulation scale, and (ii) do they yield similar spatial distributions, i.e. do the same sites tend to be frequently occupied in both the simulations and the data. For the latter question, we used correlations between site-specific simulated and predicted occupancies (averaged over time, and, for simulated data, over replicate simulations). Note that these correlations cannot be expected to exceed some limits, due to the inherent stochasticity of the simulated metapopulation. This inherent stochasticity is given by how 35-yr samples of each particular replicate simulation is correlated to the mean over all simulations - a range of values that we computed in each case.

Short-term simulations, starting with the observed data in 1983, produced global average species occupancies similar to the real data (Table S4). However, the simulations tended to predict slightly lower metapopulation occupancies than observed for *D. magna* (range from 0.11 to 0.13 instead of 0.15) and slightly higher for *D. longispina* (from 0.11 to 0.15 instead of 0.13) and *D. pulex* (from 0.06 to 0.09 instead of 0.05). The correlation between observed and simulated site-specific occupancies (averaged over 35 years) were similar to those expected based on the fundamental niches (i.e. not taking into account the initial state of the system) for *D. longispina*, slightly better for *D. magna* and slightly worse for *D. pulex* (Table S4). These correlations are, however, lower than would be expected from the stochastic variation between replicate simulations (i.e. correlation between one particular simulation and the grand mean). In the simulations, interspecific interactions did not have a large effect in general. Removing them yielded slightly lower metapopulation occupancies for all species without affecting the predicted patterns of site-specific occupancy (Table S4). Distance-limited dispersal did, however, have a detectable effect: a mean-field colonization model not accounting for distances between sites increased the expected metapopulation occupancy of *D. magna* and *D. pulex*, but reduced it for *D. longispina*, making it closer to the observed occupancies for *D. magna* and *D. longispina* but slightly further for *D. pulex*. Predictions of site-specific occupancy *r*(*x*_*dj*_, *x*_.*j*_) were improved for all species and fell within the range expected from replicate simulations (Table S4). Short-term simulations were mainly useful when initialized with the observed 1983 distribution, although that distribution was underestimated due to imperfect detection. As expected, starting with a different initial occupancy (0.2, random) tended to increase or decrease simulated occupancy depending on whether 0.2 was higher or smaller than the observed occupancy in 1983, and to weaken predictions of spatial distributions (correlations of site-specific occupancy).

Long-term simulations started with uniformly distributed occupancies that were either set to 0.2 for all species or used the distributions observed in spring 1983. Results were identical (Table S6), which shows that the memory of the initial distribution is lost over the long-term. All occupancy distributions visually converged at 2,500 years, and definitive extinctions were avoided thanks to the small offset. The offset led to slightly higher occupancies and a slight improvement in the correlations of the spatialized model (compare Tab. S5/S6). Simulations in the real landscape yielded similar results to the simulation in homogeneous landscapes (i.e. realized niches), although site-specific probabilities of occupancy were lower, and occupancy patterns were somewhat blurred in the environmental space (compare Fig. 5 and Fig. 6). Interestingly, competition had almost no effect on the site-specific occupancies, which were more (but still weakly) affected by positive interactions (See Fig. 6C). The simulations predicted a quasi-stationary metapopulation occupancy similar to that observed in the data for *D. magna*, disregarding whether dispersal limits were considered or not. The occupancy of *D. longispina* was also overall well predicted, although slightly higher than observed. Interestingly, expected occupancy for that species was higher when dispersal limitation was considered. Lastly, the long-term predicted occupancy of *D. pulex* was considerably higher than what we observed (ca. 0.15 instead of 0.05, Table S5/S6).

**Figure 6.**
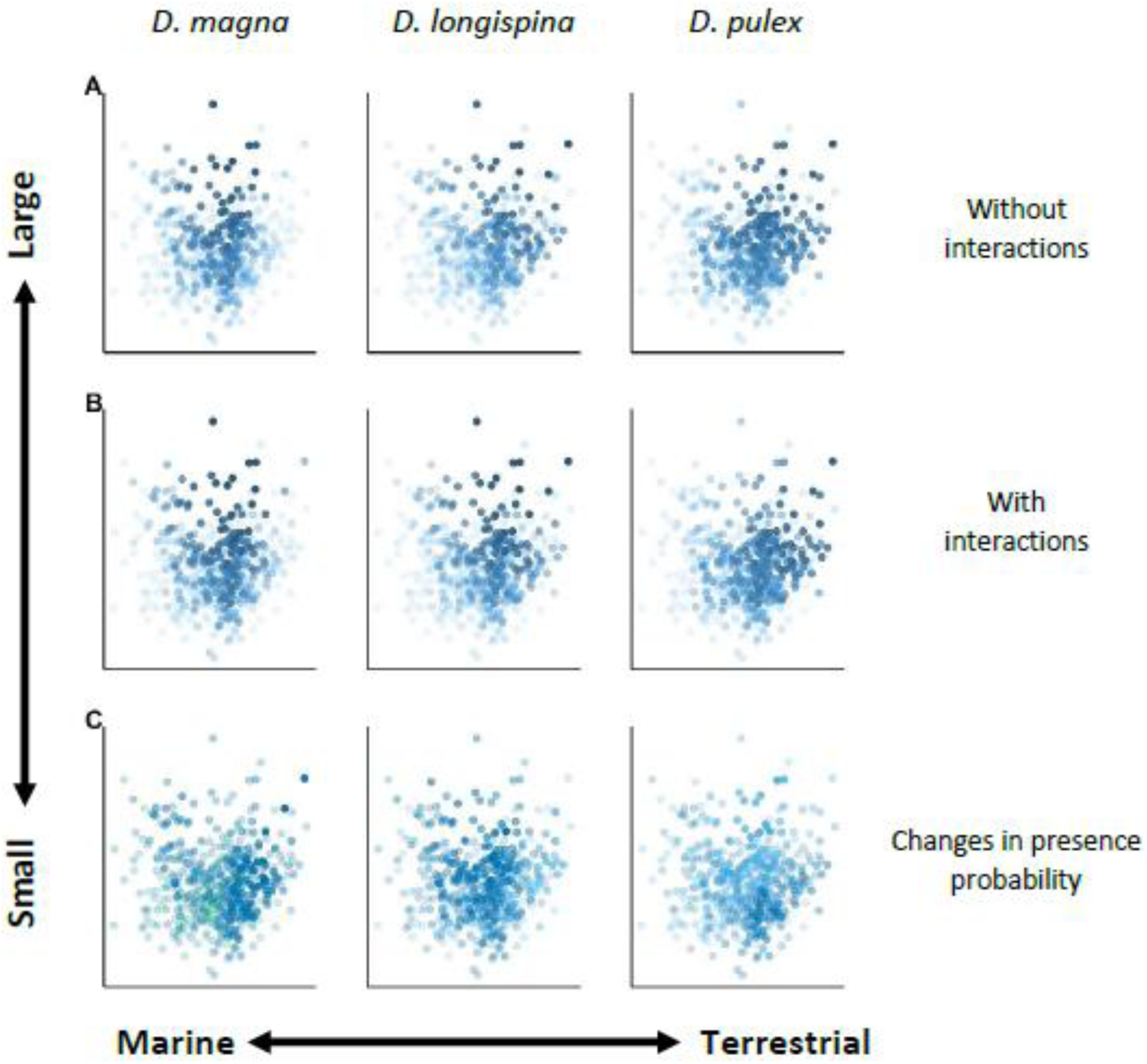
Site-specific probability of occupancy (similar to Fig. 5) based on simulations in the “true” landscape, i.e. including environmental heterogeneity and corrected for imperfect detection. In such simulations, extinctions of the whole metapopulations are very rare, and the color scale represents the average occupancy of sites among all simulations where the metapopulation was not extinct. Contrary to Fig. 5, both scales (transparency and color) represent the probability of occupancy.

The simulation models behaved quite deterministically with respect to the spatial distribution of the species, i.e. site-specific occupancy in any particular 35-yr simulated series was highly correlated with the quasi-stationary distribution (*r*(*x*_*ij*_, *x*_.*j*_) close to 0.9). In comparison, observed site-specific occupancy was poorly correlated with quasi-stationary expectations (around 0.40, 0.35 and -0.05, for *D. magna, D. longispina*, and *D. pulex*, respectively, Table S6). The impact of interspecific interactions was relatively moderate overall, as neither expected occupancy nor the patterns of site-specific occupancy were strongly modified when interactions were set to zero. However, the models were sensitive to the removal of distance effects on colonization, which affected expected metapopulation occupancy only moderately, but made the system less deterministic in two ways: (i) metapopulation occupancy varied over a larger range among replicates, and (ii) site-specific occupancy patterns were more variable across replicates (*r*(*x*_*ij*_, *x*_.*j*_) between 0.4 and 0.8 see Table S6). Nonetheless, quasi-stationary site-specific occupancy ended up predicting the data better in the model without distance-dependent colonization than in the model with it. The correlations between the two were 0.48 and 0.55 for *D. magna* and *D. longispina*, respectively, close to the maximum correlation possible based on the stochastic variation of the model and lower, though still positive at 0.21, for *D. pulex* (see Table S6).

## Discussion

### Parallel, highly seasonal, and fast-turnover extinction-colonization dynamics

In all three species, the estimated demographic rates (in an average pond, with no interactions) showed a marked difference between summer and winter, with colonization rates being much higher during summer than winter, and extinctions mostly occurring during the winter. The resulting net increase in predicted occupancy during summer and net decrease during winter corroborates with data and previous speculations based on the observation that dispersal stages (resting eggs) mainly disperse when pools fall dry in summer and the eggs are exposed to wind (Altermatt 2008, Altermatt and Ebert 2008). Low water levels in the rockpool shortly before it dries up also attract birds that feed on the animals and disperse resting eggs that survive through their gut passage (Figuerola and Green 2002, Cuhra 2019). In contrast, ponds are filled with water and may be covered by snow and ice in winter, rendering dispersal less likely. Furthermore, the presence of active (i.e. non-dormant) populations during summer can also contribute to higher colonization rates. Over short distances, planktonic *Daphnia* can be washed into neighboring pools during strong rains. On the other hand, extinction may be high during the winter because of harsh weather. Although larger resting egg banks likely help a population survive winter, some populations may not produce sufficient amounts of ephippia before the winter, so then go extinct. Furthermore, fall and winter storms increase the probability of pools being washed out by wave action or heavy rains, possibly eradicating entire populations. Lastly, predatory fish (mainly the three-spined stickleback, *Gasterosteus aculeatus*) may contribute to extinction by invading pools close to the sea, although it is unclear if this is linked to specific periods during the year.

### Similar distance-limited dispersal but different colonization rates

The estimated dispersal parameters suggest that most colonization events originate from nearby sources (median 15-20 m), with a similar distance decay for all three species. These short colonization distances make sense given the dispersal mechanisms (see previous paragraph) and correspond with previous estimates (Pajunen and Pajunen 2003). Our distance-based model is a simplification because it ignores local topography (e.g. dispersal barriers, local dendritic network, e.g. Carrara et al. 2012, Seymour et al. 2015), and may not capture rare events of long-distance dispersal. Although long-distance dispersal may constitute a tiny fraction of total colonization and thus have little influence on the fit of the dispersal kernel, it may be important for the long-term fate of the metacommunity, particularly by allowing colonization of islands that were formerly uninhabited by a given species. Yet the striking agreement among the three species and the relatively narrow credibility intervals of the estimates support the idea that the three species share the same basic, passive, dispersal mode at short distances.

At this stage, one may note that although dispersal kernels are similar among species, their basal colonization rates as well as their sensitivity to local environmental conditions are different. As the passive dispersal of ephippia seems similar, differences in colonization success may rather depend on the number and hatching success of these eggs. *D. magna* has been shown to produce more resting eggs than *D. longispina* (Zumbrunn 2009), but it has been suggested that the dispersal ability of *D. longispina* ephippia may be better than both *D. pulex*, and *D. magna*, with *D. magna* ephippia having the lowest dispersal capacity of the three species (Hanski and Ranta 1983). Environmental control can also influence successful colonization: *D. magna* cannot settle in ponds with a pH below 6.4, with experimental elevation of pH resulting in an increase of colonization rate (Ebert et al. 2013). In our model, estimates of environmental effects on colonization do not seem very large (compared to those on extinction rates); however, unfavorable environments may already be captured by the model through their effect on extinctions. Indeed, if the model predicts that populations will go extinct before the next season, the model has little information to predict whether colonization in these sites differs from the average. In addition, we cannot exclude that we may have missed potentially important environmental variables nor that some of the recorded variables have non-linear effects on colonization rates. For example, local topography determines water connection with nearby pools in a way that is not captured by our distance-based colonization function. Apparent facilitation among species on colonization may also reflect the action of such non-recorded variables (see below).

### Species-specific effects of environment on local extinctions determine fundamental niches at the metapopulation scale

In contrast to colonization rates, extinction rates seemed very sensitive to environmental parameters, although this sensitivity differed among species. A practical way to express species-specific preferences is to represent their fundamental niche, which we defined, in a metapopulation context as the set of site types, defined by environmental variables, that, if generalized across the entire metapopulation, allows persistence. Because persistence also depends on the density and spatial arrangement of habitat patches, this definition of fundamental niche only applies to a particular landscape and requires simulations that preserve the landscape characteristics. Site types that belong to the fundamental niche in our system may not do so in another system, where, for example, colonization rates may be lower because habitat patches are more distant from each other. However, although niche limits in the environmental space may expand or contract depending on patch density, the relative suitability of different site types for a species has little reason to change unless the whole environment differs. Also, different species can be compared as long as they inhabit the same landscape, as is the case here.

In our case, it turns out that the two small-bodied species (*D. pulex* and *D. longispina*) present quite similar, overlapping niches compared to the fundamental niche of the larger-bodied *D. magna*. This pattern appears to be driven mainly by species-specific sensitivities of extinction rates to both environmental axes (PC1 and PC2). The species’ position in the environmental space defined by our model strongly resembles the ordination given in Pajunen & Pajunen (2007). For example, small sites with strong marine influence are unsuitable for all three species. Furthermore, *D. magna* is known to be the most brackish water tolerant, whereas the other two are known as freshwater species (Ranta 1979, Pajunen and Pajunen 2007). It is particularly interesting that although *D. pulex* and *D. longispina* have similar fundamental niches, our model suggests that the two species have opposite seasonal patterns (i.e. summer vs. winter) of extinction rate-dependency on environmental conditions. This may indicate a form of temporal niche partitioning, that is, in a given site type, each species tends to thrive in a different season.

### Weak niche partitioning at the landscape scale despite strong competitive interactions at the local scale

Our model predicts very strong competition effects on extinction rates for all species that may or may not be symmetric, depending on species pair and season. Patterns of strong local competition, including competition effects on extinction, have previously been reported for this system (Bengtsson 1989, 1991, 1993, Hugueny et al. 2007, Zumbrunn 2009). The particularly strong effect (tenfold increase in extinction rates) of *D. pulex* and *D. longispina* on each other (in summer) is not unexpected given their similar life-histories. Species-specific parasites may also influence this dynamic, although the parasites of *D. pulex* and *D. longispina* are not much studied (Ebert et al. 2001, Zumbrunn 2009), but ongoing research suggests that they are much less parasitized than *D. magna* (D. Ebert, unpublished).

Competition may restrict the realized niches (as opposed to the fundamental ones) if it results in asymmetrical effects in some part of the ecological space. This can arise from the joint effects of environment and competition; for example, in an intermediate-sized terrestrial habitat ((−1,0) in the PC plane) where both species co-occur, extinction probability in summer is about 0.10 for *D. pulex*, and 0.26 for *D. longispina*; while it is 0.61 for *D. pulex*, and 0.04 for *D. longispina* in a more marine habitat ((1,0) in the PC plane). As a result, some habitats of the first type are removed from *D. longispina*’s niche, and some habitats of the second type from *D. pulex*’s niche, which generally coincides with the observed occupancy patterns of the two species and suggests that our model, although it simplifies matters by ignoring interactions between habitat and competition, still has the capacity to fit habitat-dependent impacts of species on each other.

At this point we are left with an apparent paradox: the two ingredients required for spatial niche partitioning, i.e., different species habitat preferences and strong local competitive interactions among species, are present; however, they do not seem to scale up to produce a strong pattern of spatial niche partitioning at the metapopulation scale. Indeed, species do not clearly segregate into different parts of the ecological space (i.e. subsets of sites) as previously supposed (Pajunen and Pajunen 2003), as we see from the fact that, globally, realized niches differ little from fundamental ones and strongly overlap between species. This may stem from several reasons: First, competitive effects on local extinction are strong but mostly symmetrical, so that in habitats broadly tolerated by two species, they may often locally exclude each other, but without a systematic advantage to one, so both species persist in this habitat type and realized niches are not or hardly reduced compared to fundamental ones. Second, the expected occupancy of the species is not very high (around 20 % of all sites), and competition only acts on doubly-occupied sites, which are rather infrequent. Third, the impact of species on each other’s extinction rates seems to be compensated by positive impacts on colonization (see below). Fourth, the high turnover of the three species in concert with habitat heterogeneity in space strongly reduces effective competitive effects at the regional scale, where only the competitive effects on *D. magna* are visible (Fig. 6C), which may explain why it was seen as a bad competitor in previous studies (Hanski and Ranta 1983).

### Apparent facilitation on colonization rates

In contrast to the negative impact observed on persistence, the presence of one *Daphnia* species in late summer tends to increase the site’s probability of being colonized by other *Daphnia* species before the next spring (i.e. winter colonization rate). The mechanism of interspecific facilitation for *Daphnia* colonization is hard to imagine, given that dispersal is passive, hatching of ephippia not known to depend on intra-or interspecific cues, and hatchling survival is probably affected by competition. However, any unmeasured factor or mechanism that makes some pools better habitats for all *Daphnia* species can results in such apparent facilitation. For instance, the absence of predators may favor colonization for all species, although predators seem not to be a major ecological factor in this system (but see Ranta and Espo 1989). Likewise, field topography may play an unrecognized role: pools located downstream in a cascade of connected pools may receive more propagules and are usually occupied more often, and colonized by new species more often than average, making it likely that a species is already present when another colonizes the site. As such specificities (and other potential effects of unmeasured variables or non-linear measured ones) are not present in our environmental variables or the current model, they will result in apparent positive interactions, potentially masking true negative interactions. Similarly, the interspecific impacts we observed on extinction may also have been underestimated because of positive correlations driven by overall lifestyle similarity and common dependency on water regimes.

Overall, the net effect of interspecific interactions - both positive and negative, apparent or real - on distribution patterns was moderate: realized niches did not differ much from fundamental ones (even slightly larger in total for *D. magna* and *D. longispina*) and still largely overlapped among the three species. While long-term relative occupancy of species in each habitat type differed, with different species being quantitatively dominant depending on the site type, few sites can be considered “private” niches, i.e. sites tolerated exclusively by only one species, confirming previous studies by Ranta (1979) and Pajunen & Pajunen (2007), which suggested that niche differences are small.

### Niche-based prediction of species distribution and the role of distance-limited dispersal

Simulations in the heterogeneous landscape based on our initial observed distributions well-reflected the observed occupancies over the 35 years’ study period, but did not predict site-specific occupancies (i.e. spatial distribution) better than the estimated niche alone. Surprisingly, predictions for *D. magna* and *D. longispina* aligned with observations better when dispersal limitations were not considered. For *D. pulex*, expected occupancy was better predicted by including dispersal limitation. When pool occupancy was randomly initialized, the correlation between simulated and real data was weaker, since less information was provided, this being particularly strong for *D. pulex*. In this regard, conclusions about dispersal kernels appear somewhat paradoxical. Indeed, the dispersal kernels themselves appear well estimated (i.e. small CI, biologically realistic and consistent with previous estimation), but simulations using these kernels generally performed worse than without, except for *D. pulex*. Long-distance colonization events, because of their apparent rarity, may explain this paradox, as these events have only a very small impact on the kernel estimation but may be important for predicting the future of the system (e.g., to repopulate isolated parts of the area that have gone extinct previously).

### The long-term behavior of the Daphnia metacommunity

Overall, the long-term projections presented the same characteristics as the short-term simulations. Again, the non-spatialized model predicted patterns of occupancies and spatial distributions quite well, showing that two species, *D. magna* and *D. longispina* would keep stable metapopulation occupancies, close to what was observed, but that *D. pulex* would reach a much higher occupancy than currently observed. Based on niche, *D. pulex* has the largest number of suitable ponds compared to the other species, yet it occupies only a small fraction of these. It is possible that *D. pulex* is slowly expanding across the archipelago, but too slowly to leave a clear signal in the 35-year series. Islands in the Tvärminne archipelago undergo modifications with time, becoming increasingly terrestrial because of post-glacial land uplift (∼30 cm/100 years), colonization and growth by trees and other plants. Such habitats are known to be more favorable to *D. pulex*, the most “continental” of the three species. On the other hand, uplifting creates new habitats as pools emerge from the sea.

### Limits of the approach

Our approach works well at disentangling spatial, environmental and interaction effects while accounting for imperfect detection. It allows proposing a mechanistic viewpoint on species dynamics in a metacommunity context and uses simulation to project dynamics under various assumptions. It enables us to see, for example, how local interactions scale at the metacommunity level when habitat heterogeneity, spatial repartition of habitat and limited dispersal are considered. With regard to the spatialized aspect, the choice of the exponential function was arbitrary and the colonization kernel could have been modeled with other distributions. In particular, long-distance events were most difficult to estimate well, leading to potentially unrealistic long-term projections. Also, some external colonization sources may exist. Indeed, although not all islands in the archipelago were sampled, some may contain *Daphnia* populations that act as stepping stones for colonization, but are not accounted for in the model. Adding an *ad hoc* small offset to the colonization rate would account for these limitations in the simulations, although further studies would benefit from addressing this issue directly.

We modeled interactions as pairwise coefficients between species independently of the environment. Yet, competition between species is likely to change depending on the environmental context (Chamberlain et al. 2014). In the current model, competitive exclusion outcomes depend on both environmental effects and interactions, but in an orthogonal way (i.e. there is no covariance between interactions and environmental effects). Similarly, we only consider first-order interactions, although higher order ones may exist. In addition, although our model considers environmental variables as fixed properties of ponds, rockpools in the Tvärminne archipelago evolve with time, as post-glacial uplifting raises islands from the sea. If such changes are important on intermediate time scales, environmental effects may be misestimated and misunderstood as interactions, as species presence is the only source of spatiotemporal variation in the predictors. For example, when a pond becomes suitable for several species due to an environmental change, environmental properties in the model do not change, so the model will attribute effects to other species, resulting in apparent facilitation. Lastly, detection probabilities are species specific and do not depend on the presence of other species or environmental factors, although these could affect it.

## Conclusion

Our study extends previous research on metacommunity dynamics both technically through its data analysis tools and biologically, with insights about the ecology of the system. It reveals marked seasonal effects, with the summer period being generally more favorable for the presence of *Daphnia* but also more dynamic. Our most striking observation is that the three species do not have identical ecological niches in the sense that their metapopulation dynamics are not affected by environmental parameters in the same way, and that they strongly compete with each other, but do not show a pattern of clear long-term segregation into ecologically distinct site subgroups (i.e. niche partitioning at the metacommunity scale). Local interactions do not scale up to the metacommunity scale because of the complex interplay of environmental heterogeneity, stochasticity in extinction and colonization processes, and spatially-limited dispersal relative to the distribution of habitats across the landscape. This echoes the fact that niche and neutral models are not opposed, and that considering a metacommunity as being driven by one or the other may depend on one’s perspective (i.e. the spatial scale). Dispersal limitation seems to be relatively well captured in our model, influencing metacommunity dynamic especially in terms of spatial distribution and number of occupied ponds. However, rare long-distance dispersal events are probably not well estimated, though they, like immigration from surrounding areas, may stabilize the metacommunity dynamic over the long term. A primary goal for future studies is to better integrate the different spatial and temporal scales at which dispersal occurs, so that long-term metacommunity trajectories can be extrapolated and closed-system assumptions relieved.

## Acknowledgements

We thank Jürgen Hottinger, Andrea Cabalzar, Mikko Lehto, Peter Fields, Jennifer Lohr, David Preiswerk, David Duneau, Katharina Ebert Gleb Ebert, A. Marcelino, Y. Haag, E. Haag, C. Liautard-Haag, C. Reisser, C. Molinier, E. Hürlimann, and C. Mills for help with data collection in the field. Jürgen Hottinger supported the laboratory work. Suzanne Zweizig improved the language of the manuscript. This work was supported by a grant of the Swiss National Science Foundation to DE and a PhD fellowship from the University of Montpellier to MD.

