## Supplements for "Species coexistence and overlapping distributions in a metacommunity are compatible with niche differences and competition at a local scale"

### Appendix A.

<u>Recovering Levins' equivalent *c*</u>

The original version of Levins' metapopulation model is not explicitly spatialized. Thus, all occupied sites (a fraction *p* of the whole set of sites*)* contribute equally to the propagule pools for colonization of an empty site, the propagule pressure being *c p*, where *c* gives the per-capita (i.e. by occupied site) colonization rate. This formula implies that, for a finite metapopulation of *n* sites, the contribution of each occupied site to the migrant pool is c/(n-1) excluding the focal site, assumed to be empty.

Under a spatialized version with dispersal limitation, propagule pressure for a particular site depends more strongly on close occupied sites than on distant ones. The contribution of each site to the propagule pool (arriving on a given site) is therefore weighted by a function of the distance between the focal site and the site sending migrants. Thus, *p* becomes specific to each receiving site that receives propagule pressure *c* $p_i$, where $p_i = \frac{1}{z}\sum_{i\neq j} x_j e^{-\delta\, D_{ij}}$, with $x_j$ the occupation status of site *j*, $1/\delta$ the mean colonization distance, $D_{ij}$ the Euclidean distance between sites *i* and *j*, and *z* a normalization constant.

The value of *z* is important because, if no normalization is made (i.e. $z = 1$), the contribution of each occupied site becomes $ce^{-\delta\, D_{ij}}$ ; the sum of these terms is not *c,* so *c* no longer has the same meaning as in the Levins' model (per-capita rate or maximal rate obtained when all sites are occupied except the focal one, i.e. all $x_j = 1$). We therefore chose the normalization constant *z* so that the average contribution of an occupied site to the propagule pool is c/(n-1) as in the unspatialized Levins' model. This results in

$$z = (n-1)\, E\left(e^{-\delta\, D_{ij}}\right) = (n-1)\frac{1}{n(n-1)}\sum_{i}\sum_{j\neq i} e^{-\delta\, D_{ij}} = \frac{1}{n}\sum_{i}\sum_{j\neq i} e^{-\delta\, D_{ij}}$$

**Figure S1.** Correlation circle from PCA on environnemental variables. Axes are reversed for presentation purposes.

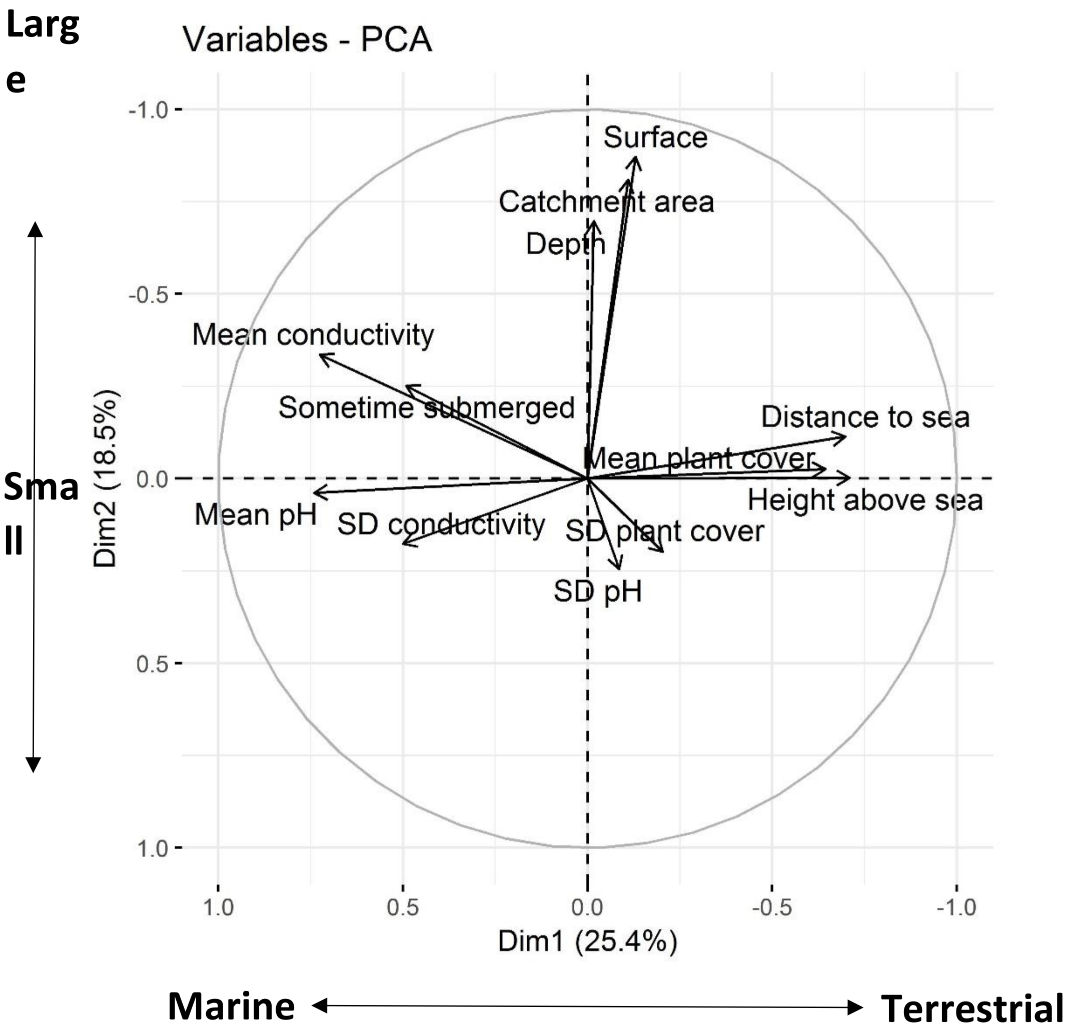

**Figure S2** Proportion of suitable sites depending on the threshold. A site is considered suitable when the proportion of non-extinct metapopulations exceeds a certain threshold at year 500, in replicate simulated homogeneous metapopulations where all sites are given the same environmental characteristics as the focal one. Thus, increasing the threshold automatically results in the inclusion of less sites on the "suitable" list.

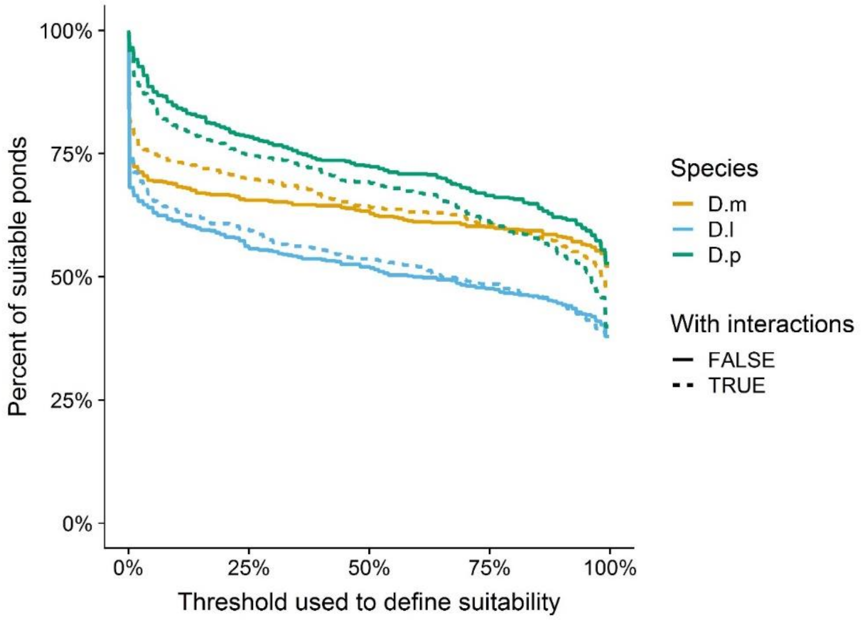

**Table S1.** Environmental variables summary. Median, mean, and SD are computed over all sites and given, for each variable, in their natural scales. Mean number of datapoints provide the mean number of measures realized by site (plus min and max). Transformations were applied before averaging measurements.

| Variable | Median | Mean | SD | Mean number of datapoints per site (min, max) | Transformation |
|---|---|---|---|---|---|
| Plant cover (%) | 0.05 | 0.15 | 0.23 | 5 [5;5] | log10(1+x) |
| Conductivity (µS) | 381.6 | 1658.3 | 2739.84 | 5.56 [5;8] | log10(x) |
| pH | 8.765 | 8.327 | 1.31 | 4.19 [4;7] | |
| Distance to sea (m) | 6.50 | 8.29 | 6.81 | | log10(x) |
| Height above sea (m) | 1.35 | 1.64 | 1.03 | | log10(x) |
| Depth (m) | 0.19 | 0.20 | 0.12 | | log10(x) |
| Surface area ($m^2$) | 1.21 | 3.30 | 11.98 | | log10(x) |
| Catchment area ($m^2$) | 8.05 | 17.54 | 39.90 | | log10(x) |
| Sometimes submerged (0/1) | 0 | 0.079 | 0.27 | | |

**Table S2.** Correlations between simulated long-term site-specific expectations (random initialization, average of 500$^{th}$-year occupancy over 500 replicates) and observed occupancies (averaged for each site over the 35 years of data). Results are obtained by simulation in homogeneous metapopulations of sites having identical environmental characteristics as focal sites. Simulations run both with interaction coefficients estimated from the data (column 1), or without (column 2).

| | With interaction coefficients | Without interaction coefficients |
|---|---|---|
| *D. magna* | 0.4802 | 0.5293 |
| *D. longispina* | 0.5631 | 0.5562 |
| *D. pulex* | 0.2293 | 0.2046 |

**Table S3.** Results of short-term simulations of metacommunity dynamics.

Each replicate simulation (*i*) started with occupancy data from Spring 1983 and ran for 35 successive years to generate a simulated time series. Replicates whose metapopulation went extinct (rare) were removed. Each year, an apparent occupancy for the summer sample was computed by randomly drawing a subset of the occupied sites based on the estimated detection probability of the species (mimicking the observation process in the observed data series). Apparent occupancies were then averaged over the 35 simulated years to obtain a site-specific apparent occupancy ($x_{ij}$ for site $j$ in replicate $i$), and the latter was averaged over all sites to get metapopulation apparent occupancy ($x_{i.}$ in replicate $i$). 500 replicate simulations were performed to estimate the expected values of apparent site-specific and metapopulation occupancy ($x_{.j}$ and $x_{..}$ respectively) over all stochastic realizations of the metacommunity model. We evaluated the fit of these expectations with the true data (*d*) based on two criteria: (1) how the observed metapopulation occupancy ($x_{d.}$) compared to the mean ($x_{..}$) and distribution of apparent metapopulation occupancy in simulations, and (2) the correlation between site-specific occupancies observed in the data ($x_{dj}$) and expected site-specific occupancies in the simulations ($x_{.j}$). For each simulation, we also evaluated the correlation with the expectation $r(x_{ij}, x_{.j})$ and provided its range to fathom the inherent stochasticity of the model. Distributions over replicated simulations are summarized as 95% ranges. The left columns (Spatialized and Interactions) indicate whether results come from simulations with or without dispersal limitation and/or species interactions.

| $\epsilon = 0$ | | *D. magna* | | *D. longispina* | | *D. pulex* | |
|---|---|---|---|---|---|---|---|
| Data ($x_{d.}$) | | 0.152 | | 0.127 | | 0.047 | |
| **Spatialized** | **Interactions** | $x_{..}$ and range of $x_{i.}$ | $r(x_{dj}, x_{.j})$: and range of $r(x_{ij}, x_{.j})$: | $x_{..}$ and range of $x_{i.}$ | $r(x_{dj}, x_{.j})$: and range of $r(x_{ij}, x_{.j})$: | $x_{..}$ and range of $x_{i.}$ | $r(x_{dj}, x_{.j})$: and range of $r(x_{ij}, x_{.j})$: |
| **X** | **X** | 0.099 (0.083-0.115) | 0.52 (0.87-0.94) | 0.153 (0.127-0.178) | 0.53 (0.81-0.88) | 0.030 (0.003-0.064) | 0.33 (0.20-0.85) |
| **X** | | 0.091 (0.076-0.106) | 0.54 (0.86-0.93) | 0.143 (0.115-0.174) | 0.56 (0.76-0.85) | 0.025 (0.003-0.061) | 0.35 (0.33-0.79) |
| | **X** | 0.131 (0.090-0.167) | 0.60 (0.51-0.68) | 0.117 (0.083-0.153) | 0.66 (0.57-0.72) | 0.079 (0.028-0.133) | 0.33 (0.285-0.59) |
| | | 0.114 (0.066-0.158) | 0.62 (0.47-0.66) | 0.108 (0.072-0.144) | 0.66 (0.52-0.70) | 0.072 (0.018-0.141) | 0.35 (0.27-0.57) |

**Table S4**. Results of short-term simulations of metacommunity dynamics, including an external colonization source ($\epsilon = 5.10^{-4}$). Legend as in Table S3. The additional left column (Initialization) indicates whether the metacommunity was initialized with Spring 83 distribution (83), or at random with probability 0.2 (R).

| $\epsilon = 5.10^{-4}$ | | | *D. magna* | | *D. longispina* | | *D. pulex* | |
|---|---|---|---|---|---|---|---|---|
| Data ($x_{d.}$) | | | 0.152 | | 0.127 | | 0.047 | |
| **Spatialized** | **Interactions** | **Initialization** | $x_{..}$ and range of $x_{i.}$ | $r(x_{dj}, x_{.j})$: and range of $r(x_{ij}, x_{.j})$: | $x_{..}$ and range of $x_{i.}$ | $r(x_{dj}, x_{.j})$: and range of $r(x_{ij}, x_{.j})$: | $x_{..}$ and range of $x_{i.}$ | $r(x_{dj}, x_{.j})$: and range of $r(x_{ij}, x_{.j})$: |
| **X** | **X** | **83** | 0.121 (0.091-0.160) | 0.54 (0.79-0.91) | 0.154 (0.130-0.179) | 0.54 (0.81-0.88) | 0.068 (0.024-0.114) | 0.14 (0.29-0.81) |
| **X** | | **83** | 0.106 (0.079-0.141) | 0.57 (0.78-0.91) | 0.146 (0.113-0.179) | 0.56 (0.76-0.85) | 0.055 (0.014-0.104) | 0.17 (0.22-0.76) |
| | **X** | **83** | 0.134 (0.094-0.173) | 0.60 (0.52-0.69) | 0.120 (0.085-0.154) | 0.66 (0.58-0.72) | 0.088 (0.036-0.146) | 0.33 (0.29-0.60) |
| | | **83** | 0.119 (0.079-0.163) | 0.62 (0.50-0.67) | 0.113 (0.075-0.152) | 0.66 (0.53-0.69) | 0.085 (0.035-0.148) | 0.33 (0.28-0.58) |
| **X** | **X** | **R** | 0.181 (0.149-0.163) | 0.43 (0.50-0.89) | 0.162 (0.133-0.152) | 0.40 (0.80-0.89) | 0.160 (0.125-0.194) | -0.036 (0.65-0.83) |
| **X** | | **R** | 0.164 (0.137-0.193) | 0.47 (0.78-0.87) | 0.150 (0.116-0.183) | 0.42 (0.69-0.82) | 0.177 (0.133-0.223) | -0.07 (0.69-0.82) |
| | **X** | **R** | 0.166 (0.118-0.216) | 0.49 (0.50-0.68) | 0.132 (0.090-0.178) | 0.55 (0.51-0.71) | 0.166 (0.116-0.218) | 0.22 (0.42-0.67) |
| | | **R** | 0.149 (0.108-0.188) | 0.52 (0.48-0.67) | 0.117 (0.078-0.162) | 0.53 (0.36-0.60) | 0.180 (0.120-0.240) | 0.19 (0.37-0.67) |

**Table S5**. Results of long-term simulations of metacommunity dynamics. Each replicate simulation (*i*) started with the Spring 1983 occupancy data and ran for 2,500 successive years to generate a simulated time series, of which only the last 35 years were considered. Occupancies were corrected for detectability as indicated in Table S3.

| $\epsilon = 0$ | | *D. magna* | | *D. longispina* | | *D. pulex* | |
|---|---|---|---|---|---|---|---|
| Data ($x_{d.}$) | | 0.152 | | 0.127 | | 0.047 | |
| **Spatialized** | **Interactions** | $x_{..}$ and range of $x_{i.}$ | $r(x_{dj}, x_{.j})$: and range of $r(x_{ij}, x_{.j})$: | $x_{..}$ and range of $x_{i.}$ | $r(x_{dj}, x_{.j})$: and range of $r(x_{ij}, x_{.j})$: | $x_{..}$ and range of $x_{i.}$ | $r(x_{dj}, x_{.j})$: and range of $r(x_{ij}, x_{.j})$: |
| **X** | **X** | 0.155 (0.124-0.194) | 0.34 (0.86-0.95) | 0.137 (0.114-0.162) | 0.28 (0.91-0.95) | 0.148 (0.103-0.191) | -0.07 (0.76-0.87) |
| **X** | | 0.141 (0.116-0.174) | 0.38 (0.86-0.93) | 0.126 (0.101-0.148) | 0.30 (0.85-0.92) | 0.177 (0.118-0.227) | -0.09 (0.77-0.89) |
| | **X** | 0.177 (0.109-0.238) | 0.49 (0.54-0.72) | 0.128 (0.062-0.184) | 0.55 (0.54-0.75) | 0.185 (0.121-0.246) | 0.21 (0.45-0.71) |
| | | 0.155 (0.124-0.194) | 0.34 (0.86-0.95) | 0.137 (0.114-0.162) | 0.28 (0.91-0.95) | 0.148 (0.103-0.191) | -0.07 (0.76-0.87) |

**Table S6**. Results of long-term (2,500y) simulations of metacommunity dynamics, including an external colonization source ($\epsilon = 5.10^{-4}$). Legend as in Table S4. The additional left column (Initialization) indicates whether the metacommunity was initialized with Spring 83 distribution (83), or randomly with probability 0.2 (R).

| $\epsilon = 5.10^{-4}$ | | | *D. magna* | | *D. longispina* | | *D. pulex* | |
|---|---|---|---|---|---|---|---|---|
| Data ($x_{d.}$) | | | 0.152 | | 0.127 | | 0.047 | |
| **Spatialized** | **Interactions** | **Initialization** | $x_{..}$ and range of $x_{i.}$ | $r(x_{dj}, x_{.j})$: and range of $r(x_{ij}, x_{.j})$: | $x_{..}$ and range of $x_{i.}$ | $r(x_{dj}, x_{.j})$: and range of $r(x_{ij}, x_{.j})$: | $x_{..}$ and range of $x_{i.}$ | $r(x_{dj}, x_{.j})$: and range of $r(x_{ij}, x_{.j})$: |
| **X** | **X** | **83** | 0.180 (0.144-0.220) | 0.39 (0.85-0.93) | 0.156 (0.121-0.194) | 0.35 (0.86-0.93) | 0.165 (0.129-0.200) | -0.04 (0.75-0.89) |
| **X** | | **83** | 0.161 (0.135-0.188) | 0.43 (0.85-0.93) | 0.146 (0.113-0.187) | 0.36 (0.79-0.88) | 0.190 (0.148-0.236) | -0.07 (0.78-0.89) |
| | **X** | **83** | 0.180 (0.120-0.239) | 0.48 (0.55-0.73) | 0.134 (0.077-0.184) | 0.55 (0.57-0.75) | 0.186 (0.126-0.251) | 0.21 (0.43-0.72) |
| | | **83** | 0.160 (0.106-0.216) | 0.52 (0.54-0.75) | 0.108 (0.042-0.167) | 0.53 (0.37-0.65) | 0.22 (0.139-0.290) | 0.20 (0.46-0.74) |
| **X** | **X** | **R** | 0.181 (0.144-0.221) | 0.39 (0.85-0.93) | 0.156 (0.122-0.193) | 0.35 (0.86-0.93) | 0.163 (0.124-0.199) | -0.04 (0.73-0.88) |
| **X** | | **R** | 0.159 (0.131-0.186) | 0.43 (0.85-0.92) | 0.145 (0.112-0.183) | 0.36 (0.79-0.89) | 0.190 (0.144-0.239) | -0.076 (0.77-0.89) |
| | **X** | **R** | 0.184 (0.144-0.246) | 0.49 (0.54-0.73) | 0.135 (0.069-0.193) | 0.56 (0.56-0.75) | 0.189 (0.121-0.253) | 0.22 (0.49-0.72) |
| | | **R** | 0.163 (0.108-0.214) | 0.53 (0.53-0.72) | 0.107 (0.045-0.166) | 0.53 (0.36-0.65) | 0.221 (0.143-0.290) | 0.19 (0.47-0.74) |
